# NF1 deficiency induces metabolic reprogramming and epithelial-mesenchymal transition in glioblastoma

**DOI:** 10.64898/2026.06.17.733017

**Authors:** Qiang Dong, Jing Shi, Hang Yin, Bo Wang, Liang Niu, Xiaoqing Wang, Junqing Dai, Qiang Li, Yawen Pan, Guoqiang Yuan

## Abstract

**Background:** Metabolic reprogramming is a common occurrence in tumor cells, where enhanced glycolysis promotes cell growth, invasion and migration. NF1 is tumor suppressor gene that downregulates the encoded neurofibromin protein. However, the effects of NF1 on energy metabolism and epithelial-mesenchymal transition (EMT) in glioblastoma multiforme (GBM), as well as the underlying molecular mechanisms, remain unclear.

**Methods:** CRISPR/Cas9 gene editing technology was employed to construct GBM cell lines with NF1 gene mutations. Metabolomics was utilized to examine the impact of NF1 on metabolic remodeling in GBM. The Seahorse XF24 extracellular flux analyzer was used to detect the effect of NF1 knockdown on glycolysis and mitochondrial oxidative phosphorylation in GBM cells. Wound healing assay and Transwell chamber assay were utilized to detect the effect of NF1 on GBM cell invasion. Orthotopic tumor model in nude mice was established to explore the role of NF1 in *vivo*. In addition, Co-IP, western blotting, and immunofluorescence were used to explore the changes of key enzymes in glycolysis and mitochondrial oxidative phosphorylation and the relationship between NF1 and MFN1.

**Results:** The expression of NF1 is decreased in glioma tissues and is significantly correlated with patient prognosis. NF1 knockdown may promote the invasion, migration, and EMT of GBM cells. At the same time, the activation of the AKT/mTOR signaling pathway promotes aerobic glycolysis in GBM cells, promotes mitochondrial division through targeted regulation of MFN1, and inhibits mitochondrial oxidative phosphorylation. NF1 deficiency promotes EMT in GBM cells by enhancing aerobic glycolysis and mitochondrial division.

**Conclusion:** NF1 deficiency promotes GBM glycolysis by activating the AKT/mTOR signaling pathway and inhibits the mitochondrial oxidative phosphorylation by regulating MFN1; NF1 deletion promotes GBM EMT by remodeling the pattern of energy metabolism.

## Introduction

Glioblastoma multiforme (GBM) is the most aggressive brain tumor in adults, with a five-year survival rate of only 5.5% [1]. Currently, the pathological grading of GBM is primarily based on morphology. However, tumors with similar tissue characteristics often exhibit distinct molecular genetic profiles, leading to significant variations in prognosis among individuals of the same grade [2]. This indicates that a patient’s molecular characteristics directly influence tumor behavior and play a critical role in guiding treatment, evaluating therapeutic efficacy, and predicting prognosis. Large-scale sequencing analyses from The Cancer Genome Atlas (TCGA) have classified GBM into four molecular subtypes: classical, mesenchymal, proneural, and neuronal. NF1 is inactivated through genetic loss or mutation in approximately 15% of glioma cases, with a mutation rate as high as 11% in GBM [3, 4]. Among the four molecular subtypes of GBM, the mesenchymal subgroup is often associated with NF1 gene deletions and mutations [5].

Neurofibromin 1 (NF1) is a tumor suppressor gene, located in region 11.2 of the long (q) arm of chromosome 17. It comprises 62 exons and encodes 2818 amino acids, with a protein molecular weight of 280 kDa [6]. The NF1 gene is expressed in all cells, with the highest expression observed in neurons, Schwann cells, glial cells, and white blood cells. Neurofibromatosis protein is encoded by the tumor suppressor gene NF1, and its deletion results in tumor susceptibility syndrome type 1 neurofibromatosis. Neurofibromin acts as a tase-activating protein of RAS (RAS-GAP). Neurofibromin is involved in a variety of cell signaling pathways, including Ras/MAPK, AKT/mTOR, ROCK/LIMK/cofilin, and cAMP/PKA pathways, and regulate cell proliferation and migration [7]. Studies have indicated that the molecular changes that occur in NF1-related gliomas of different grades are inconsistent. In low-grade gliomas, the inactivation of NF1 mutations often activates the Raf/MEK/ERK signaling pathway, which affects the immune function of tumor immune cells and promotes tumorigenesis [8, 9]. The inactivation of NF1 mutations in high-grade gliomas often activates the PI3K/ AKT/mTOR signaling pathway, which affects the mitosis, cellular energy metabolism and nerve regeneration in tumor cells, thereby promoting tumor progression. NF1 mutations promote the phosphorylation of AKT and mTOR in astrocytes, activate the AKT/mTOR signaling pathway, and inhibit the growth of astrocytes after treatment with mTOR inhibitors [10]. Currently, the ERK inhibitor (Selumetinib) has been used in phase II and III studies in NF1-related low-grade gliomas and optic pathway gliomas [11, 12]. Sokol et al. discovered that NF1 mutations result in acquired resistance to tamoxifen in invasive lobular carcinoma of the breast cancer. NF1 gene mutations occur mainly in lobular lung cancer, and NF1 mutations are more frequent in metastatic lobular lung cancer [13]. Su et al. found that the deletion of NF1 in ovarian cancer can induce the upregulation of MCL1 expression and render ovarian cancer cells anti-apoptotic through miR- 142-5p [14]. They further demonstrated that the loss of NF1 inhibited cisplatin induced apoptosis and causes ovarian cancer cells to become resistant to chemotherapy.

The epithelial-mesenchymal transition (EMT) is closely related to tumor metastasis. EMT may also induce cell dedifferentiation of non-cancer stem cells, allowing the cells to acquire self-renewal and tumor initiation capabilities, as well as tolerance to chemoradiotherapy [15]. Although a large amount of literature has studied the role of EMT in cancer, its application in cancer diagnosis and treatment is still limited, partly due to the heterogeneity and microenvironment differences in tumor tissues [16]. In fact, the invasive ability of the tumor does not originate from the tumor tissue mass but depends on the microenvironment of the specific tissue. Therefore, tumor microenvironment plays an important role in promoting the occurrence of tumor EMT. Zhang et al.[17] found that deletion of the NF1 gene can down-regulate the expression of E-cadherin, a marker on the surface of epithelial cells, and reduce the ability of cell invasion and migration by inhibiting the EMT in mice. At the same time, in neurofibromatosis type 1, decreased expression of neurofibromatosis protein also promotes EMT [18]. NF1 plays an important role in regulating EMT.

Activated oncogenes and inactivated tumor suppressors contribute to tumorigenesis by reprogramming cellular energy and metabolism toward macromolecular synthesis, a phenomenon known as metabolic reprogramming [19]. Metabolic reprogramming of tumor cells plays an important role in maintaining the growth and proliferation of tumor cells. Tumor cells in the tumor microenvironment with sufficient oxygen, tumor cells usually show an energy metabolism mode based on aerobic glycolysis, which provides energy for tumor cells and a carbon skeleton for nucleic acid synthesis [20, 21]. Tumor cell metabolism has been considered as a therapeutic hotspot for dietary and drug interventions. NF1 can regulate the GTPase activity of Ras, and RAS-activating mutations drive metabolic programming and introduce energetic and metabolic dependencies [22]. Numerous studies have explored the role of RAS mutations in tumor energy metabolism. However, the biological characteristics and energy metabolism patterns of GBM cells caused by NF1 mutations remain unclear, and the pathogenesis and microenvironment driven by genetic and epigenetic factors should be better defined. In this study, we found that the expression of NF1 in glioma patients was significantly correlated with patient prognosis. NF1 deficiency promoted aerobic glycolysis of GBM cells and inhibited oxidative phosphorylation of mitochondria. At the same time, NF1 promotes mitochondrial division through its interaction with MFN1, thereby reducing mitochondrial function, and deletion of NF1 gene enhances glycolysis in GBM cells through activation of AKT/mTOR signaling pathway and promotes the occurrence of EMT. These mechanisms provide a new research direction for the development of anti-glioma drugs and a new ideas for the treatment of glioma.

## Materials and methods

### 2.1 Patients and tissue specimens

34 glioma tissues (8 grade II,12 grade III and 14 grade IV) were collected from the Department of Neurosurgery of the Second Hospital of the Lanzhou University. The 8 normal brain tissues from patients who were diagnosed with deep brain GBM and had to undergo fistula surgery. The tissue samples were snap-frozen in liquid nitrogen and stored at -80°C until used for RNA isolation.

### 2.2 Reagents

MG-132 and 3-MA were purchased from Selleck Chemicals (Shanghai, China). The Glycolytic Rate Test kit and Mito Pressure Test Kit were purchased from Agilent Technology Co. LTD (Wilmington, USA). MitoTracker™ Green FM was purchased from Thermo Fisher Scientific (Waltham, MA, USA). Reactive Oxygen Species (ROS) Assay Kit was purchased from Beyotime Biotechnology (Shanghai, China). Bax, Bcl-2, ND1, SHDB, UQCRC2, MTCO2, ATP5A, MFN2, GLUT1, HK2, PMK2, LDH, HIF-1a and Drp1 antibodies were obtained from proteintech (Wuhan, China). AKT, p-AKT, mTOR and p-mTOR antibodies were obtained from Cell Signaling Technology. GAPDH and NF1 antibodies were obtained from Abcam (Cambridge, UK).

### 2.3 Lentiviral transfection

U87 cells (cell density 2×10^5^/ml) were cultured in 6-well plate. Adds the amount of NF1 knocked down virus with an MOI of 5. After infection in a cell incubator for 12 h, replace the solution containing the virus was replaced complete culture medium. Puromycin at a final concentration of 8 μg/ml was added to virus-infected cells several times to screen for stable NF1 knockout cell lines.

### 2.4 Generation of NF1-Knockout U251 Cells with the CRISPR/Cas9 System

NF1-KO U251 cells were constructed using the CRISPR/Cas9 system targeting the human NF1 exon 2 regions to ensure an effective single-guide RNA. The sgRNA sequence: gRNA#1: TTCCCAGGGAA GGGCTCGTTTGG; gRNA#2: CCAGAGAAATGAGTTTGTCTGGG.

### 2.5 Mitochondrial Morphology Imaging

To observe mitochondrial morphology, Mito Tracker Green CMXRos (Invitrogen) was used to mark the mitochondria of U251/U87 cells. Mito Tracker was diluted with basal medium to a concentration of 20 nM and incubated at 37 °C for 30 min in the dark and fixed with 4% paraformaldehyde, then observed and analyzed under a fluorescence microscope.

### 2.6 Mitochondrial Stress Test Assay and Glycolysis Stress Test Assay

U251/U87 cells were seeded on XF24 microplates 20000/well and incubated overnight. The mitochondrial Stress Test Assay and Glycolysis Stress Test Assay were used to detect cellular mitochondrial oxidative (OCR) phosphorylation and glycolysis capacity (ECR) using an the XF24 Seahorse Biosciences Extracellular Flux Analyzer (Seahorse Bioscience, 102238-100). Five replicates were performed for each experimental group for analysis.

### 2.7 Wound healing assay

GBM cells were plated into a 6-well plate. When the cells reached a confluence of 95%, they were gently and slowly scratched with a new 200 ml pipette tip. Cells migrating was monitored and measured using a bright-field microscope after culturing the cells for 48 h in serum-free medium. The experiments were repeated three times.

### 2.8 Invasion and transwell assay

Approximately 1×10^6^ cells in 100 μl serum-free medium were added to Matrigel coated transwell chamber, and 600μl of DMEM with 10% FBS was added to the lower transwell compartment. After 24 hours of cultivation, the cells at the bottom of the small chamber were fixed with a 4% polymerization formaldehyde, and 0.1% crystalline purple staining was cleaned with PBS. Images of cells were captured using a bright-field microscope. Cell invasion assays were performed as described above, except for cell culture inserts coated with Matrigel (BD Biosciences).

### 2.9 Western blot

After centrifugation, total protein was extracted with RIPA buffer, and the concentration was examined via BCA protein analysis kit. SDS-PAGE was used to separate the protein and transferred onto polyvinylidene fluoride (PVDF) membranes. The membranes were incubated with primary antibodies for overnight at 4 °C, and incubated with appropriate peroxidase-conjugated secondary antibodies for 1.5 h at room temperature, and then visualized by enhanced chemiluminescence using image-Quant LAS 500 system.

### 2.10 ROS measurements

To evaluate intracellular ROS level, NF1 knockdown U251/U87 cells were incubated with 10 μM DCFH-DA (Solarbio, CA1410) for 20 min at 37°C, then washed with serum free medium for three times and imaged using Olympus fluorescence microscope (BX53). The cells were then digested with trypsin and resuspended. ROS levels were analyzed using flow cytometry (BD FACSCantoTM low cytometry, USA).

### 2.11 Transmission electron microscopy

Cells collected by trypsinization were fixed with 2.5% glutaraldehyde, followed by 1%OsO4. After dehydration, the thin sections were stained with uranyl acetate and observed under a transmission electron microscope (JEM-1230, JEOL, Japan).

### 2.12 Animals experiments

Four-week-old female BALB/c nude mice were bred under SPF conditions and were randomly assigned to each group (eight mice per group). A total of 1 × 10^7^ U87 cells were injected into the right basal ganglia of mice (seam coronal and sagittal suture node as the center, to the right side to open 2 mm, to 1 mm forward as a vaccination point). Four weeks later, brains were harvested and analyzed. The Lanzhou University of Second Hospital Ethical Committee approved all the animal experiments.

### 2.13 Histology and immunohistochemistry analysis

Four weeks after tumor inoculation, all animals were sacrificed, and their subcutaneous tumor were excised and fixed in 4% paraformaldehyde and embedded in paraffin. Sections (5 μm thick) were stained with hematoxylin and eosin (HE) and immunohistochemical staining.

### 2.14 Statistical analysis

The data were analyzed using SPSS22.0 software. All the figures were performed using GraphPad Prism software. The student’s t-test and One-way Analyses of Variance (ANOVA) with a Tukey’s post-hoc test was used to assess group differences. Error bars represent the standard error of the mean (SEM). A P value <0.05 was statistically significant.

## 3. Results

### 3.1 NF1 is lowly expressed in human glioma

To analyze NF1 expression in tumor tissues, we used data from The Cancer Genome Atlas (TCGA) and found that NF1 mRNA levels in GBM tissue were lower than those in normal brain tissue. Interestingly, in certain tumor types, such as cholangiocarcinoma, gastric adenocarcinoma, head and neck squamous cell carcinoma, liver hepatocellular carcinoma, lung adenocarcinoma, and lung squamous cell carcinoma, NF1 mRNA expression was significantly elevated compared to that in normal tissues (Figure 1A). To further verify the expression of NF1 mRNA, qPCR was performed on 34 gliomas of different grades and 8 normal brain tissues. Studies have shown that the expression of NF1 mRNA in glioma tissues is significantly decreased, and with the increase of glioma grade, the expression level of NF1 is lower and lower (Figure 1D). Next, we used the Human Protein Atlas and CGGA databases to analyze the expression of NF1 at different levels again, and the results showed that, consistent with the previous experimental results, the higher the level, the lower the expression of NF1 (Figure 1B and C). In addition, we used the CGGA database to analyze the relationship between NF1 and patient survival prognosis of patients. The result showed that Lower NF1 expression was associated with shorter survival times and poorer prognosis in patients with glioma (p < 0.0001) and GBM patients (p = 0.013) (Figure 1E and F).

**Figure 1.**
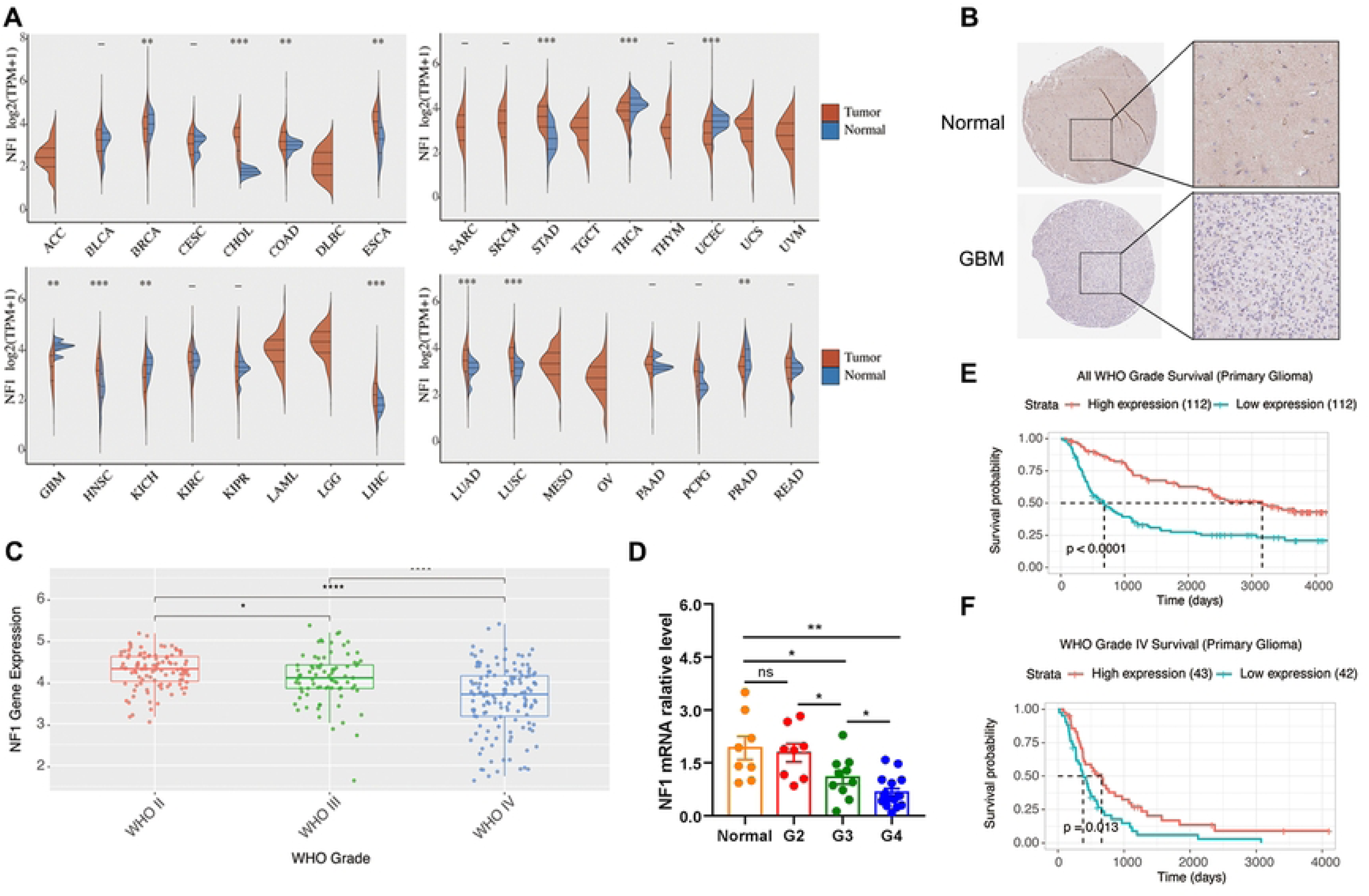
NF1 expression is reduced in glioma tissues and associated with patient prognosis. (A) Violin plot shows the expression of NF1 mRNA in the TCGA database; (B) Human Protein Atlas database analyzed NF1 protein expression between GBM and normal. (C) Box plot shows the expression of NF1 mRNA in different grades of glioma in the CGGA database; (D) qPCR detected the expression of NF1 mRNA in different grades of glioma tissues; (E and F) Kaplan-Meier curves were used to analyze the relationship between NF1 and the prognosis of glioma and grade IV glioma patients in the CGGA database.

### 3.2 NF1 deficiency promotes the invasion and transwell of GBM cells

To investigate the effects of NF1 gene knockout on GBM cell invasion and migration, we transfected U87 cells with an shNF1 lentiviral vector to establish a stable NF1 knockdown cell line. Scratch wound healing assays revealed that cells in the NF1 knockdown group migrated faster than those in the control group, with a wound closure rate of approximately 60% compared with 45% in the control group after 24 hours of incubation in a 6-well plate (Figures 2F and G). Similar results were observed in the Transwell chamber assay, where a greater number of cells invaded and migrated through the chamber in the NF1 knockdown group compared to the control group (Figures 2A-E). These findings suggest that NF1 gene deletion promotes GBM cell invasion and migration.

**Figure 2.**
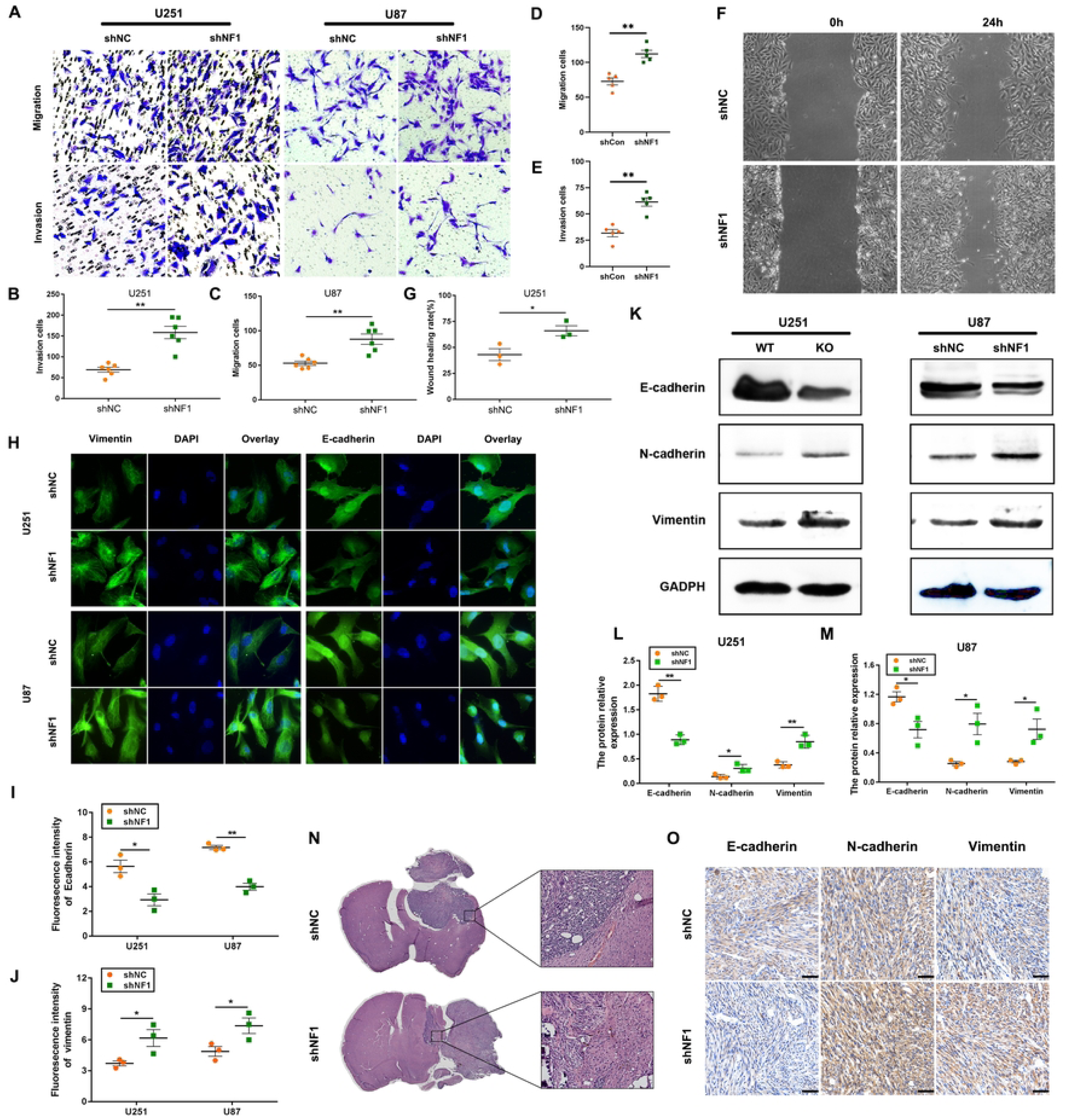
NF1 gene knockdown promotes GBM cell invasion and migration. (A) Transwell chamber assay was used to examine the effect of NF1 knockout on the invasion and migration of GBM cells; (B-E) Statistical analysis of Figure A. (F) The cell scratch healing assay was used to detect the effect of NF1 knockout on U251 cell migration; (G) Statistical analysis of Figure F. (H) Immunofluorescence was used to detect the changes in E-cadherin and Vimentin protein expression after NF1 gene knockdown in U251 and U87 cells; (I-J) Image J was used to analyze the changes in the fluorescence intensity of the Vimentin and E-cadherin protein in Figure H; (K) Western blotting was used to detect the expression changes of EMT-related proteins E-cadherin, N-cadherin and Vimentin after NF1 gene knockdown in U251 and U87 cells; (L-M) The relative expression of E-cadherin, N-cadherin, and Vimentin proteins in U87 and U251 cells in Figure I was analyzed; (N) HE was used to observe the pathological morphology of orthotopic transplanted tumors in nude mice; (O) E-cadherin, N-cadherin, and Vimentin determined by immunohistochemistry in Orthotopic xenograft tumor model. *: P < 0.05, **: P < 0.01.

EMT is a morphological process in which epithelial cells acquire mesenchymal characteristics. EMT is closely associated with tumor progression and has been a major focus of cancer research. To investigate the impact of NF1 deficiency on EMT, we used immunofluorescence and western blotting to detect changes in EMT-related molecules. Immunofluorescence analysis revealed a significant decrease in the expression of the epithelial marker E-cadherin and a marked increase in the mesenchymal marker Vimentin in U251 and U87 cells with NF1 knockdown (Figures 2H, I, and J). Western blot analysis confirmed these findings, showing a significant reduction in E-cadherin expression and an increase in N-cadherin and Vimentin expression in NF1 deficiency cells (Figures 2K, L, and M). Additionally, hematoxylin and eosin (HE) staining of orthotopic transplanted tumors showed that the boundary between the tumor and normal tissue was indistinct, with clear infiltrative tumor growth and pronounced cellular atypia in the NF1 loss group (Figure 2N). Immunohistochemical analysis of tumor tissues further corroborated these results, demonstrating reduced E-cadherin and increased N-cadherin and Vimentin expression in NF1 knockdown tumors compared to controls (Figure 2O). These data indicate that NF1 knockdown promoted EMT in GBM cells.

### 3.3 NF1 Regulates the Glycolytic Capacity of GBM

To investigate the impact of NF1 gene defects on cellular energy metabolism, we analyzed metabolites using ultra-high-performance liquid chromatography coupled with tandem mass spectrometry (LC-MS/MS). A Metware Database (MWDB) was developed based on standard products for the qualitative analysis of the mass spectrometry data. Our results showed that 20 out of the 57 energy metabolites analyzed displayed significant differences compared to the control group. KEGG pathway analysis indicated that these altered metabolites were primarily involved in pyrimidine metabolism, vancomycin resistance, vitamin B metabolism, amino sugar and nucleotide sugar metabolism, as well as monomycin metabolism (Figures 3A). Notably, among the differentially regulated metabolites, glucose-6-phosphate and lactate levels were significantly elevated in the glycolytic pathway. Additionally, there was a negative correlation between NF1 expression and the lactic acid content. These findings suggest that NF1 knockout may affect energy metabolism in GBM cells, particularly through its impact on the glycolytic pathway, potentially contributing to tumor initiation and progression.

**Figure 3.**
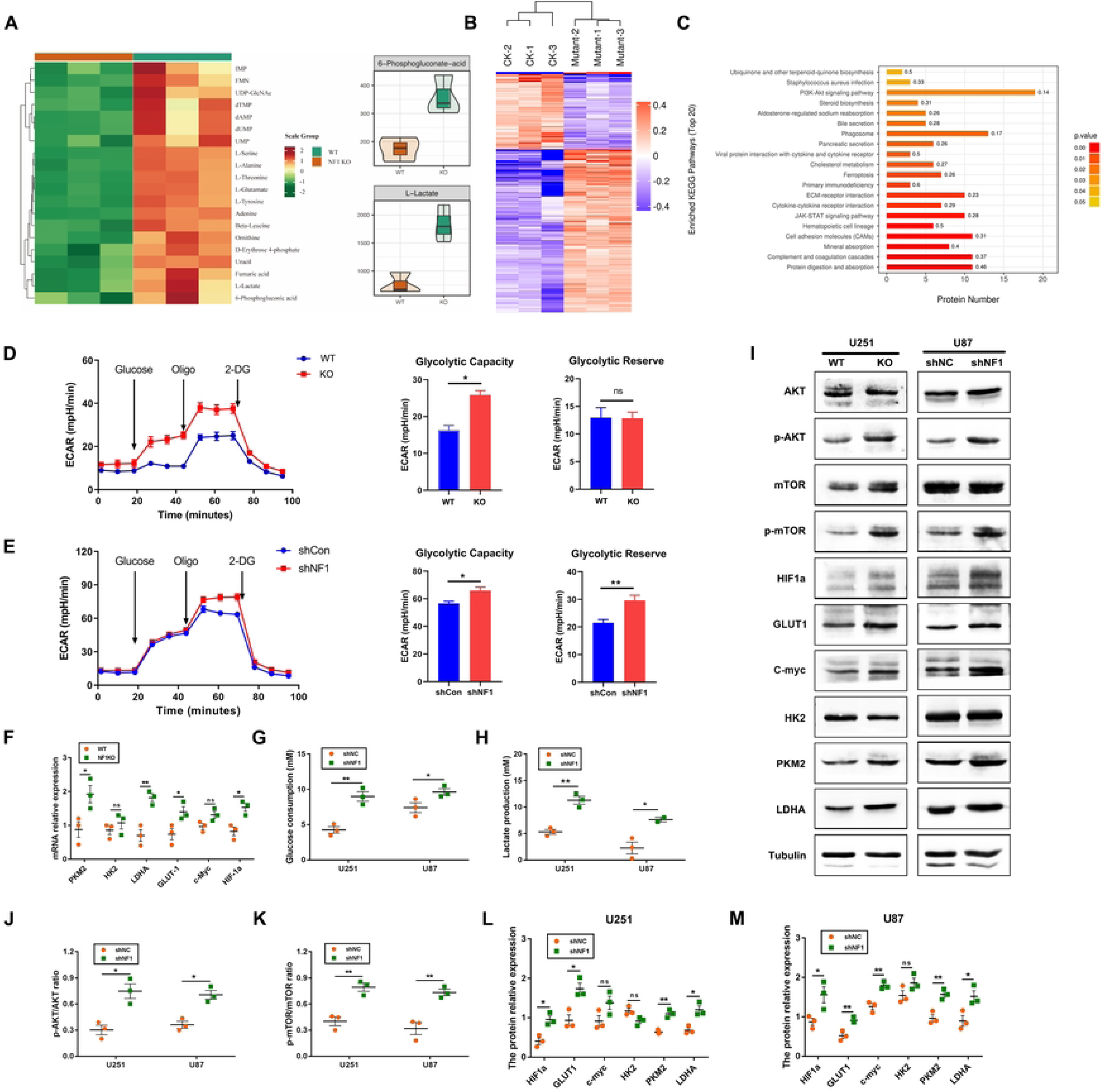
Knockdown of NF1 gene promotes aerobic glycolysis in U251 and U87 cells. (A) The heat map represents the differential metabolic products of GBM cells after NF1 knockout, and the violin plot represents the contents of glucose 6-phosphate and lactate; (B) Proteomics was used to detect the changes of differential proteins between GBM cells and normal cells after NF1 knockdown.; (C) Differential protein signaling pathway enrichment; (D-E) The Seahorse XF24 extracellular flux analyzer was used to detect the effect of NF1 gene knockout on glycolytic capacity in U251 and U87 cells; (I) WB detection was used to detect the expression of proteins related to the AKT/mTOR signaling pathway and glycolysis-related proteins (GLUT-1, PKM2, HK2, LDHA, C-Myc, HIF1a); (F) qRT-PCR was used to detect the changes in the expression of mRNA of glycolysis pathway-related molecules in U251 cells with NF1 gene knockout; (G) A glucose detection kit was used to detect the effect of glucose consumption in the cells of the NF1 gene knockout group and the control group; (H) A lactate detection kit was used to detect the effect of lactate production in the cell culture medium of the NF1 gene knockout group and the control group; (J-M) The relative expression of glycolysis pathway-related proteins in U87 and U251 cells in Figure I was analyzed. *: P < 0.05, **: P < 0.01.

Next, after transfection with lentivirus, we collected the cell culture medium in the logarithmic growth phase and tested the glucose and lactic acid content in the culture medium using a glucose and lactic acid detection kit. The results showed that compared with the control group, the consumption of glucose and the production of lactic acid in the medium of the NF1 knockdown group was significantly increased, with statistical significance (p < 0.05) (Figure 3G and H). At the same time, we used Seahorse XF24 extracellular flow analyzer to examine the effect of NF1 gene knockout on glycolysis capacity. The results showed that transfection of lentivirus knockdown NF1 in U251 and U87 cells significantly increased the glycolytic capacity and glycolytic storage capacity of the cells compared with the control group (Figure 3D and E). We further detected the changes in mRNA levels of molecules related to the glycolysis pathway by qPCR, and the results showed that the mRNA expression levels of PKM2, LDHA, C-myc, and HIF1a in U251 cells with NF1 gene knockout were significantly increased compared to the control group (Figure 3F). At the same time, WB was used to detect the changes of glycolytic pathway-related protein molecules, and we found that the results were consistent with the results of energy metabolomics and qPCR. The expressions of PKM2, LDHA, Glut-1 and HIF1a was increased in U251 and U87 cells with NF1 knockdown. Interestingly, HK2 and C-myc did not differ in the expression of U251 protein upon NF1 knockdown, whereas C-myc expression was significantly reduced in U87 cells with NF1 knockdown (Figure 3I). These results suggest that NF1 gene knockdown may activate the glycolytic pathway and promote lactic acid production. The molecular mechanism by which NF1 deficiency in promoting glycolysis was investigated. The effects of NF1 gene knockout on the protein expression profiles of wild-type GBM cells were analyzed using the iTRAQ technique, revealing 379 up-regulated proteins and 172 down-regulated proteins. Additionally, KEGG enrichment analysis indicated the activation of the PI3K/AKT signaling pathway (Figure 3B and C). Studies have shown that the AKT/ mTOR/HIF-1a pathway plays an important role in glycolysis. We further used WB to detect changes in the AKT/ mTOR/HIF-1a signaling pathway. The results showed that after the deletion of NF1, the phosphorylation levels of AKT and mTOR and the expression of HIF-1a were significantly enhanced, activating the AKT/mTOR/HIF-1a signaling pathway (Figure 3I-M), which was consistent with the results of previous studies. In conclusion, NF1 gene deficiency may activate the AKT/ mTOR/HIF-1a signaling pathway and enhance the glycolytic ability of GBM cells.

### 3.4 NF1 deficiency increases mitochondrial fragmentation in Glioma Cells

Mitochondrial morphology plays an important role in the regulation of cellular metabolism. In this study, we analyzed the effect of NF1 loss on mitochondrial morphology using the MitroTracker and transmission electron microscopy. As shown in the figure 4A, confocal microscopy revealed that in NF1knockdown U251 and U87 cells, the mitochondria appeared smaller, more fragmented, and significantly shortened in length. In contrast, the control cells exhibited a higher number of large, filamentous mitochondria with compact cristae. Consistent with MitroTracker staining, the mitochondria were smaller and vacuolated in the knockdown cells as observed by transmission electron microscopy (Figure 4B). Therefore, we hypothesized that mitochondrial fragmentation leads to mitochondrial metabolic dysfunction.

**Figure 4.**
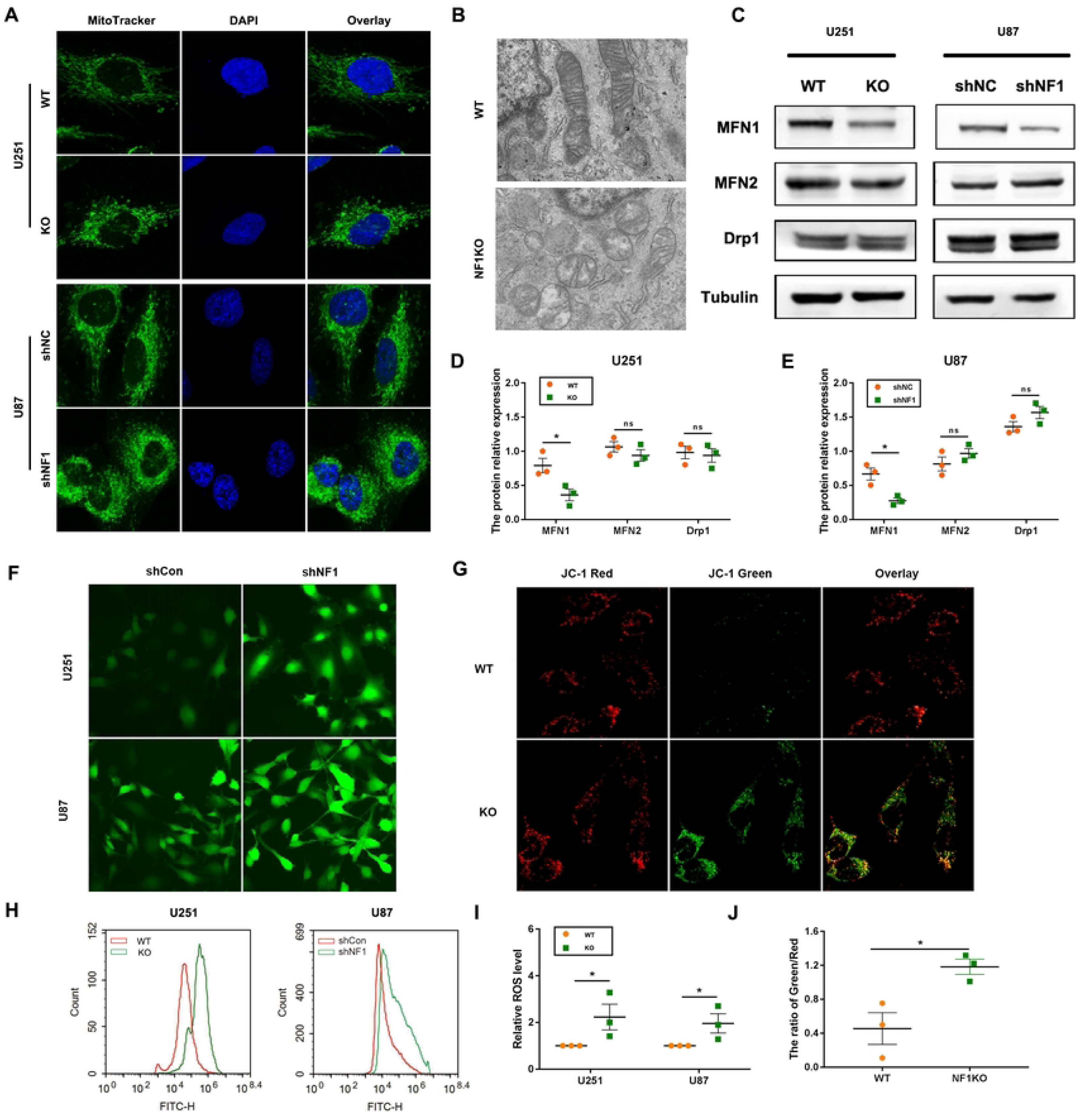
NF1 gene knockout promotes mitochondrial fission in GBM cells. (A) Mitochondria in the NF1 knockdown group and the control group were stained with MitoTracker dye, and changes in mitochondrial morphology were observed under a confocal microscope; (B) Changes in mitochondrial morphology in NF1 knockout U251 cells were observed under a transmission electron microscope; (C) Western blot was used to detect changes in mitochondrial dynamics-related proteins (MFN1, MFN2, Drp1); (D-E) Quantitative analysis of the protein bands in Figure C; (F) DCFH-DA reactive oxygen species detection reagent was used to stain NF1-knockdown U251 and U87 cells, and the changes in intracellular reactive oxygen species were observed under a fluorescence microscope; (H) The DCFH-DA kit was used to detect the changes in intracellular reactive oxygen species under a flow cytometer; (G) The JC-1 detection kit was used to observe the changes in mitochondrial membrane potential in NF1-knockdown U251 cells under a fluorescence microscope; (I) Image J was used to quantitatively analyze the fluorescence intensity in Figure F; (J) Image J was used to analyze the ratio of green and red fluorescence in Figure G. *: P < 0.05, **: P < 0.01.

To further understand the mechanism by which NF1 regulates mitochondrial dynamics, we examined the expression of proteins related to mitochondrial dynamics, including MFN1, MFN2 and Drp1. In NF1 loss GBM cells, western blotting results showed that the expression of MFN1 was significantly decreased, whereas the expressions of MFN2 and Drp1 was not significantly changed (Figure 4C, D, and E). In summary, we found that NF1 loss affects the changes in mitochondrial morphology and function. The molecular mechanism may involve NF1 exerting its role by targeting MFN1 regulation. The rapid proliferation of cancer cells results in a high reliance on ATP for energy. Compared to normal cells, the clearance mechanism of intracellular reactive oxygen species (ROS) in tumor cells has changed. In tumor cells, some ROS can stimulate the growth of tumor cells. Therefore, we employed the reactive oxygen species detection kit to detect the changes in ROS in NF1 knockdown cells. As shown in the figure, the reactive oxygen species in U251 and U87 cells in the NF1 knockdown group were significantly increased (Figure 4F and I). At the same time, we used flow cytometry to detect the changes in intracellular reactive oxygen species, and we found that the results were consistent with the results of the kit detection, and the reactive oxygen species in the knockdown group were significantly increased (Figure 4H). When generating energy, mitochondria store electrochemical potential energy in the mitochondrial inner membrane, on both sides of the inner membrane. If the concentration of protons and other ions is asymmetrically distributed, the mitochondrial membrane potential is formed. A normal mitochondrial membrane potential is a prerequisite for maintaining mitochondrial oxidative phosphorylation and adenosine triphosphate production. The stability of the mitochondrial membrane potential is conducive to maintaining the normal physiological function of cells. Therefore, we used the JC-1 kit to detect the changes in mitochondrial membrane potential of U251/U87 cells. The results showed that after NF1 gene knockdown, the mitochondrial membrane potential was significantly reduced (Figure 4G and J). These results suggest that NF1 is involved in the regulation of mitochondrial function and cellular energy metabolism.

### 3.5 Loss of NF1 inhibits mitochondrial oxidative phosphorylation

The Warburg effect is characterized by abnormal metabolic phenomena that enhance glycolysis and reduce mitochondrial oxidative phosphorylation, leading to significant differences between cancer cells and normal cells and affecting tumor progression. Previous studies have shown that deletion of NF1 gene enhances glycolysis in GBM cells. Next, we further investigated its effect on mitochondrial oxidative phosphorylation to explore whether deletion of the NF1 gene reshapes the energy metabolism pattern of glioma cells. First, we used a Seahorse XF24 extracellular flow analyzer to examine the effect of NF1 gene knockdown on mitochondrial oxidative phosphorylation. Compared with the control group, the mutation of NF1 gene significantly increased the glycolytic capacity and glycolytic storage capacity of GBM cells. From the oxygen consumption rate curve, we found that compared with the control group, the basic and maximum mitochondrial respiratory capacity of U251 and U87 cells was decreased after transfection with lectin virus knocked down NF1, and ATP production was significantly reduced (Figure 5A and B). At the same time, the content of ATP in their cells was detected by ATP detection kit. The results showed that the ATP content of the cells in the knockdown group was significantly reduced compared with that in the control group (Figure 5C). Mitochondrial ATP is mainly by the mitochondrial oxidative phosphorylation of ATP release energy, due to the generation of ATP is reduced, so we further tested the coenzyme molecular expression associated with mitochondrial energy production, according to the results, the NF1 knock on low cells, mitochondria coenzyme I (ND1), The expressions of mitochondrial Coenzyme II (SDHB) and mitochondrial Coenzyme III (UQCRC2) were significantly decreased. However, mitochondrial coenzyme IV (MTCO2) and mitochondrial Coenzyme V (ATP5A) protein expressions did not change significantly (Figures 5D-F). In summary, NF1 gene knockout reduced mitochondrial oxidative phosphorylation, reshaped the energy metabolism pattern of GBM cells, and promoted the occurrence of the Warburg effect.

**Figure 5.**
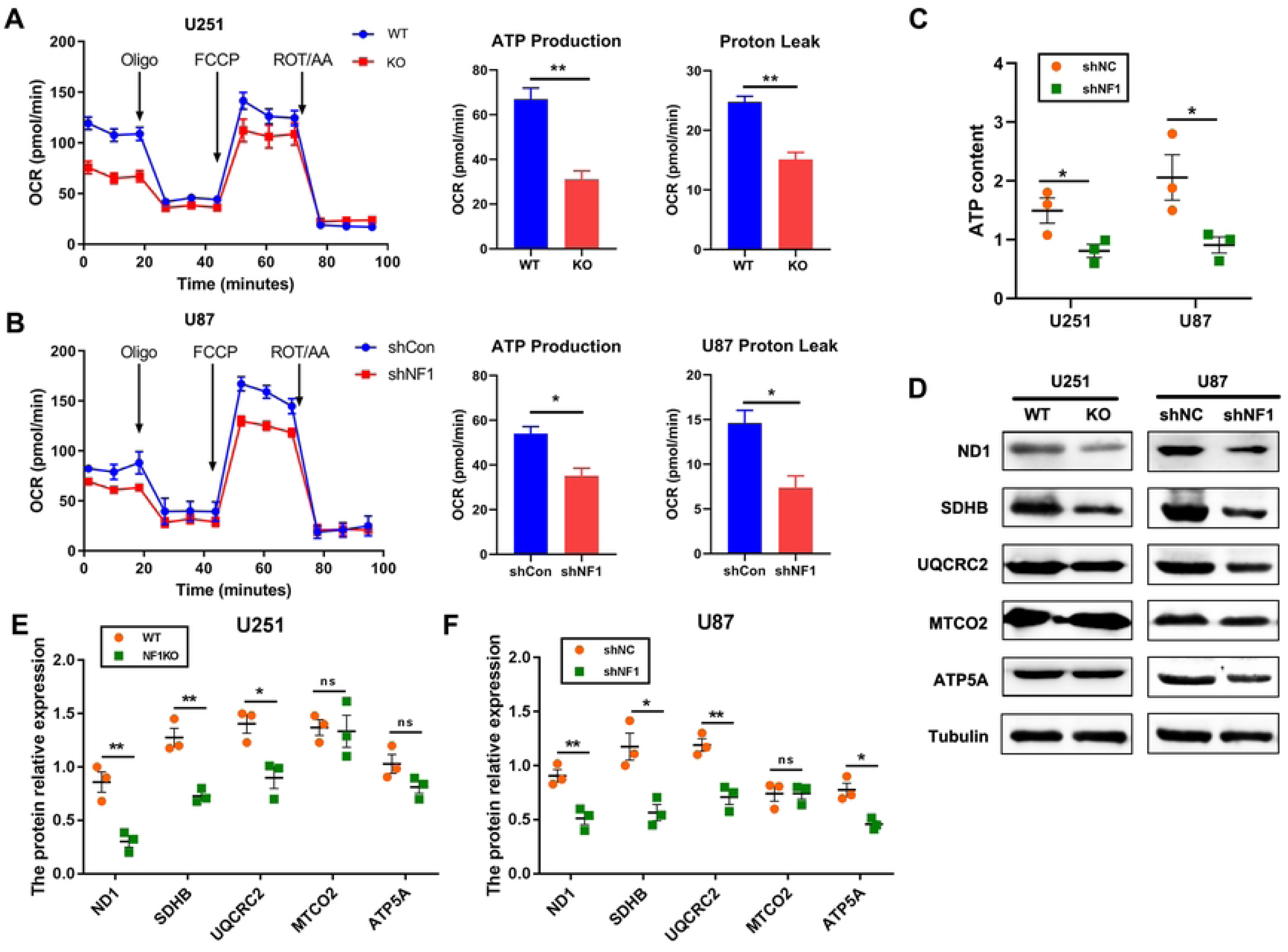
NF1 gene knockout inhibits mitochondrial oxidative phosphorylation in GBM cells. (A) The Seahorse XF24 extracellular flux analyzer detected the effect of NF1 gene knockout in U251 cells on mitochondrial oxidative phosphorylation; (B) The Seahorse XF24 extracellular flux analyzer detected the effect of NF1 gene knockdown in U87 cells on mitochondrial oxidative phosphorylation; (C) The ATP detection kit was used to detect the changes in ATP content in NF1 knockdown U251 and U87 cells; (D) The expression of mitochondrial oxidative phosphorylation-related proteins (ND1, SDHB, UQCRC2, MTCO2, ATP5A) was detected by WB detection; (E-F) The relative expression of mitochondrial complex enzymes in Figure D in U87 and U251 cells was analyzed. *: P < 0.05, **: P < 0.01.

### 3.6 NF1 promotes mitochondrial fission by regulating MFN1

We found that NF1 knockdown promoted mitochondrial division, and the expression of MFN1 was also reduced. We hypothesized that NF1 promoted mitochondrial division by regulating MFN1 expression. Further, by analyzing the mRNA data of GEPIA and CGGA databases, we found that MFN1 was positively correlated with NF1, with correlation coefficients of 0.45 and 0.277, respectively (Figure 6C and D). This is consistent with our WB results. We then observed the intracellular spatial localization of NF1, MFN1, and mitochondria using immunofluorescence (Figures 6A and B). In U251 and U87 cells, double staining of NF1 with MFN1 and mitochondria showed that NF1 colocalized with mitochondria and that NF1 colocalized with MFN1. These results suggest that NF1 can recruit mitochondria to regulate MFN1expression. To further verify this conclusion, we used co-immunoprecipitation to verify the direct interaction between NF1 and MFN1. As shown in the figure, MFN1 protein was found in the NF1 immunoprecipitation complex, and both NF1 protein were present in the MFN1 immunoprecipitation complex, which further confirmed the direct interaction between NF1 and MFN1 (Figure 6E).

**Figure 6.**
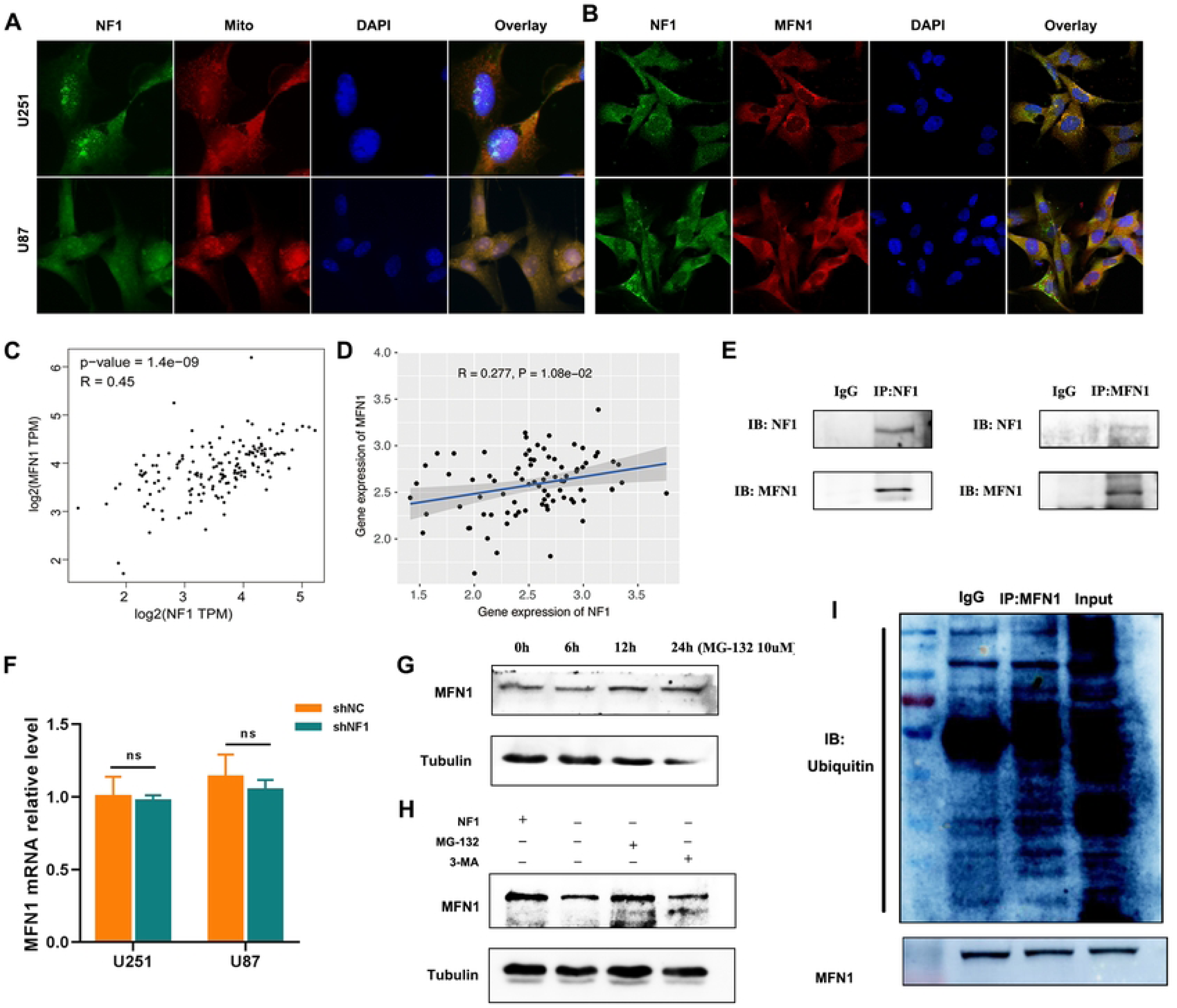
NF1 deficiency promotes mitochondrial fission by regulating MFN1. (A) Immunofluorescence double staining of NF1 and MitoTracker in U87 and U251 cells to detect the spatial relationship between NF1 and mitochondria; (B) Immunofluorescence double staining of NF1 and MFN1 in U87 and U251 cells to detect the spatial relationship between NF1 and MFN1 protein expression; (C) GEPIA databases analyzed the relationship between NF1 and MFN1; (D) CGGA databases analyzed the relationship between NF1 and MFN1; (E) Protein immunoprecipitation was used to detect the interaction between NF1 and MFN1 proteins; (F) RT-qPCR was used to detect changes in MFN1 mRNA levels in NF1-knocked U251 and U87 cells; (G) The autophagy-lysosome inhibitor 3-MA and the ubiquitin-proteasome inhibitor MG-132 were added to NF1-knocked U251 cells, and the changes in MFN1 protein expression were detected by WB; (H) NF1-knocked U251 cells were treated with MG-132 for different time periods to detect changes in MFN1 protein expression; (I) Protein co-immunoprecipitation was used to detect the interaction between NFN1 and ubiquitin. *: P < 0.05, **: P < 0.01.

The expression of MFN1 protein was significantly reduced after NF1 knockdown; however, the pathway of MFN1 degradation remains unclear. We first used qPCR to detect the changes in MFN1 transcription levels and found that NF1 gene knockdown did not affect the changes in MFN1 mRNA levels (Figure 6F). Therefore, we added autophagy lysosome inhibitor 3-MA and proteasome inhibitor MG-132 (10 uM) to NF1 knockdown U251 cells, and the results showed that MFN1 expression in NF1 knockdown group added MG-132 was significantly higher than that in the 3-MA and control group. The expression of MFN1 increased with prolonged MG-132 was treatment (Figure 6H). Next, we used co-immunoprecipitation to verify whether MFN1 interacts with ubiquitin. As shown in the figure, MFN1 interacted with total ubiquitin (Figure 6I). Taken together, these results show that MFN1 is degraded via the ubiquitin proteasome pathway.

## Discussion

GBM is the most malignant brain tumor with poor prognosis, high mortality and significant invasiveness, with a five-year survival rate of only 5.5% [2]. Despite advancements in emerging technologies such as electric field therapy and immunotherapy, the current standard treatment protocol for GBM remains maximal safe resection followed by radiotherapy and temozolomide chemotherapy. Nevertheless, even with aggressive treatment, the median survival time for GBM patients is only 12 to 18 months [1, 23]. Presently, the pathological grade of GBM mainly relies on its pathological morphology. However, GBM with identical or similar tissue characteristics can exhibit considerable variability in clinical outcomes, resulting in large prognostic differences between individuals with the same grade. GBM can be classified into proneuronal, interstitial, neuronal, and classical subtypes at the molecular level, based on variations in the expression of GBM molecules, gene mutations, and DNA methylation levels. The prognoses of patients with these distinct subtypes exhibit significant differences [24, 25]. This indicates that the molecular characteristics of patients directly influence the biological behavior of tumors and are crucial for guiding treatment, assessing therapeutic efficacy, and predicting prognosis. This study used data from the TCGA, GEPIA and GCCA databases, together with in vivo and in vitro experiments, to show that NF1 expression is reduced in GBM tissues. Furthermore, lower NF1 expression correlates with poorer prognosis in GBM patients. Energy metabolomics analysis further revealed that NF1 knockdown promotes glycolysis in glioma cells while inhibiting mitochondrial oxidative phosphorylation. Protein immunoprecipitation assays showed that NF1 directly interacted with MFN1 to promote mitochondrial division. NF1 deficiency reshaped the energy metabolism pattern of GBM cells and promoted GBM progression. Deletion of NF1 gene promotes EMT in GBM cells by activating the AKT/mTOR signaling pathway to promote glycolysis and by targeting MFN1 to promote mitochondrial division.

In this study, we analyzed NF1 gene expression in various cancers using the TCGA database. Notably, NF1 expression was found to be negatively correlated with the malignancy level in certain cancers, including glioma, breast cancer, thyroid cancer, and endometrial cancer, and positively correlated with malignancy in cancers such as cholangiocarcinoma, gastric adenocarcinoma, head and neck squamous cell carcinoma, liver hepatocellular carcinoma, and lung adenocarcinoma and squamous cell carcinoma. These findings suggest that NF1 may play a dual role in cancer progression, depending on the tumor type. We also analyzed the relationship between NF1 expression and patient prognosis using the CGGA database. Our analysis revealed that higher NF1 expression is associated with better patient prognosis and is inversely correlated with glioma grade. Thus, NF1 may serve as a prognostic marker for GBM patients. Consistent with previous reports, NF1 expression was found to be significantly reduced in optic glioma. Solga et al. demonstrated that transgenic mice lacking NF1 in the central nervous system exhibited multiple glial cell defects, including a developmental defect leading to the proliferation of reactive astrocytes and increased glial progenitor cell proliferation in the adult brain, resulting in optic nerve enlargement [26]. As a result, all mutated optic nerves developed hyperplastic lesions, which eventually progressed into optic glioma. Qiao et al. found that NF1 expression was significantly reduced in ovarian cancer and that this reduction was strongly associated with lymph node metastasis in ovarian cancer [27]. Meanwhile, NF1 can also serve as a prognostic marker for ovarian cancer, with lower NF1 expression being associated with reduced five-year survival rates. Mutations in the NF1 gene lead to the downregulation of the neurofibromin protein it encodes, which in turn activates the RAS signaling pathways (HRAS, NRAS, and KRAS-RAF-MAPK), promoting tumor proliferation and metastasis. In this study, we found that NF1 knockdown enhanced glioma cell invasion and migration compared with the control group.

The EMT plays a key role in cancer cell migration, invasion, and metastasis. It may also induce the dedifferentiation of non-cancer stem cells, allowing them to acquire self-renewal, tumor-initiating properties, and resistance to radiotherapy and chemotherapy [28, 29]. Using GBM cells with NF1 knockdown, we observed morphological changes, including a more elongated shape. In our previous experiments, we found that NF1 deficiency significantly enhanced the metastatic and invasive capabilities of U251 and U87 cells. Meanwhile, we detected changes in EMT markers, with a significant decrease in epithelial markers and an increase in mesenchymal markers. Zhang et al [30] found that NF1 gene can down-regulate the expression of E-cadherin, a marker on the surface of epithelial cells, in mice’s blood-abundant cells, and reduce cell invasion and migration ability by inhibiting EMT. This is consistent with the results of our study. In neurofibromatosis type 1, reduced expression of neurofibromatosis protein also promotes EMT. At the same time, some scholars have reported that decreased expression of NF1 in ovarian cancer promotes lymph node metastasis of ovarian cancer [31]. Therefore, NF1 gene is closely related to the occurrence of EMT, but the specific mechanism is still unclear. Therefore, understanding the driving mechanisms of EMT is important for identifying new targets to prevent the diffuse infiltration of GBM cells. Changes in the tumor microenvironment present significant challenges for cancer treatment. Many tumor tissues exhibit higher glucose uptake than adjacent normal tissues [32, 33]. Therefore, targeting tumor cell glycolysis and inhibiting the mitochondrial respiratory chain has become a critical approach for treating malignant tumors. This study uses NF1 knockdown U251 cells to investigate energy metabolism, which has important clinical implications for the treatment of NF1 mutated gliomas. AKT, a serine/threonine kinase, phosphorylates downstream targets and plays a key role in cancer growth and metabolism. Research has indicated that AKT activity promotes the Warburg effect. The PI3K/AKT/mTOR signaling pathway has also been implicated in the upregulation of HIF-1α [34]. It upregulates the transcription of glucose transporters and almost all glycolytic enzymes, such as HK2, PKM2 and LDHA [35]. In this study, using the Seahorse XF24 extracellular flux analyzer, we found that glycolytic activity was significantly increased in the NF1 knockdown group, suggesting that GBM cells with NF1 knockdown are better adapted to the hypoxic tumor microenvironment. Additionally, NF1 knockdown promoted Akt and mTOR phosphorylation, thereby activating the AKT/mTOR signaling pathway. This activation was accompanied by a marked increase in the levels of Glut-1, PKM2, and LDHA, which are key downstream molecules of HIF-1α. These findings indicate that NF1 deletion-induced AKT/mTOR pathway activation plays a crucial role in elevating glycolytic enzyme levels (Glut-1, PKM2, and LDHA) and enhancing aerobic glycolysis in GBM cells. As previously reported, enhanced glycolysis can drive tumor cell invasion, migration, and EMT. Thus, NF1 deletion may promote EMT through the activation of the AKT/mTOR pathway, leading to increased glycolysis in GBM cells.

Increasing evidence indicates that mitochondria play a critical role in tumorigenesis and tumor progression. As a dynamic network, mitochondria constantly alter their structure to meet cellular energy demands and to adapt to the extracellular environment. Abnormal expression of dynamin-related proteins, including dynamin 1-like (DNM1L), mitofusin-1 (MFN1), and mitofusin-2 (MFN2), has been observed in glioma, lung cancer, colon cancer, and breast cancer [36, 37]. Recent studies have demonstrated that lung cancer exhibits excessive mitochondrial fission and impaired fusion owing to an imbalance in DNM1L/MFN expression, which is crucial for cell cycle progression [38]. Additionally, significant upregulation of DNM1L has been shown to promote breast cancer cell metastasis by enhancing mitochondrial division. Increased mitochondrial fragmentation facilitates the survival of hepatoma cells by promoting autophagy and inhibiting mitochondria-dependent apoptosis [39, 40]. In cancer cells with dysfunctional mitochondria, elevated intracellular ROS levels may increase the effectiveness of drug therapies. In this study, we found that NF1 knockdown promoted mitochondrial division and decreased the mitochondrial OCR levels in glioma cells. The expression of mitochondrial fusion protein MFN1 decreased, resulting in the imbalance of DNM1L/MFN expression, mitochondrial ATP production, decreased membrane potential in U251 and U87 cells, and increased ROS production increased, triggering mitochondrial apoptosis. Deletion of the NF1 gene promotes the occurrence of EMT in GBM cells, which may be achieved by enhancing cellular glycolysis and promoting mitochondrial division.

MFN1 plays a crucial role in the maintaining of mitochondrial stability. The absence of MFN1 promotes mitochondrial division [41, 42]. Through protein co-immunoprecipitation, we identified a direct interaction between NF1 and MFN1. Following NF1 gene knockout, MFN1 protein expression was significantly decreased, demonstrating a positive correlation between NF1 and MFN1 expression. However, NF1 knockdown did not alter NF1 mRNA levels, suggesting that MFN1 underwent post-translational modification. Previous studies have identified two primary pathways for protein degradation: the autophagic lysosomal pathway and the ubiquitin-proteasome pathway. To investigate this in glioblastoma (GBM) cells, we treated the cells with the autophagic lysosome inhibitor 3-MA and the ubiquitin-proteasome inhibitor MG-132. Following MG-132 treatment, MFN1 protein expression increased, suggesting that MFN1 was degraded via the ubiquitin-proteasome pathway. Furthermore, protein immunoprecipitation confirmed that MFN1 binding to ubiquitin promotes its degradation.

In summary, by analyzing multiple databases and clinical glioma tissues, we observed that NF1 expression was significantly reduced in glioma patients and closely correlated with patient prognosis. NF1 deficiency promoted aerobic glycolysis in GBM cells and inhibited mitochondrial oxidative phosphorylation. Additionally, NF1 regulates mitochondrial division by interacting with MFN1, thereby impairing mitochondrial function. The loss of NF1 enhances glycolysis and promotes epithelial-mesenchymal transition (EMT) in GBM cells by activating the AKT/mTOR signaling pathway. Simultaneously, NF1 deletion promoted EMT by regulating MFN1 to increase mitochondrial division (Figure 7). These findings offer new insights into the development of anti-glioma therapies and provide novel approaches to glioma treatment.

**Figure 7.**
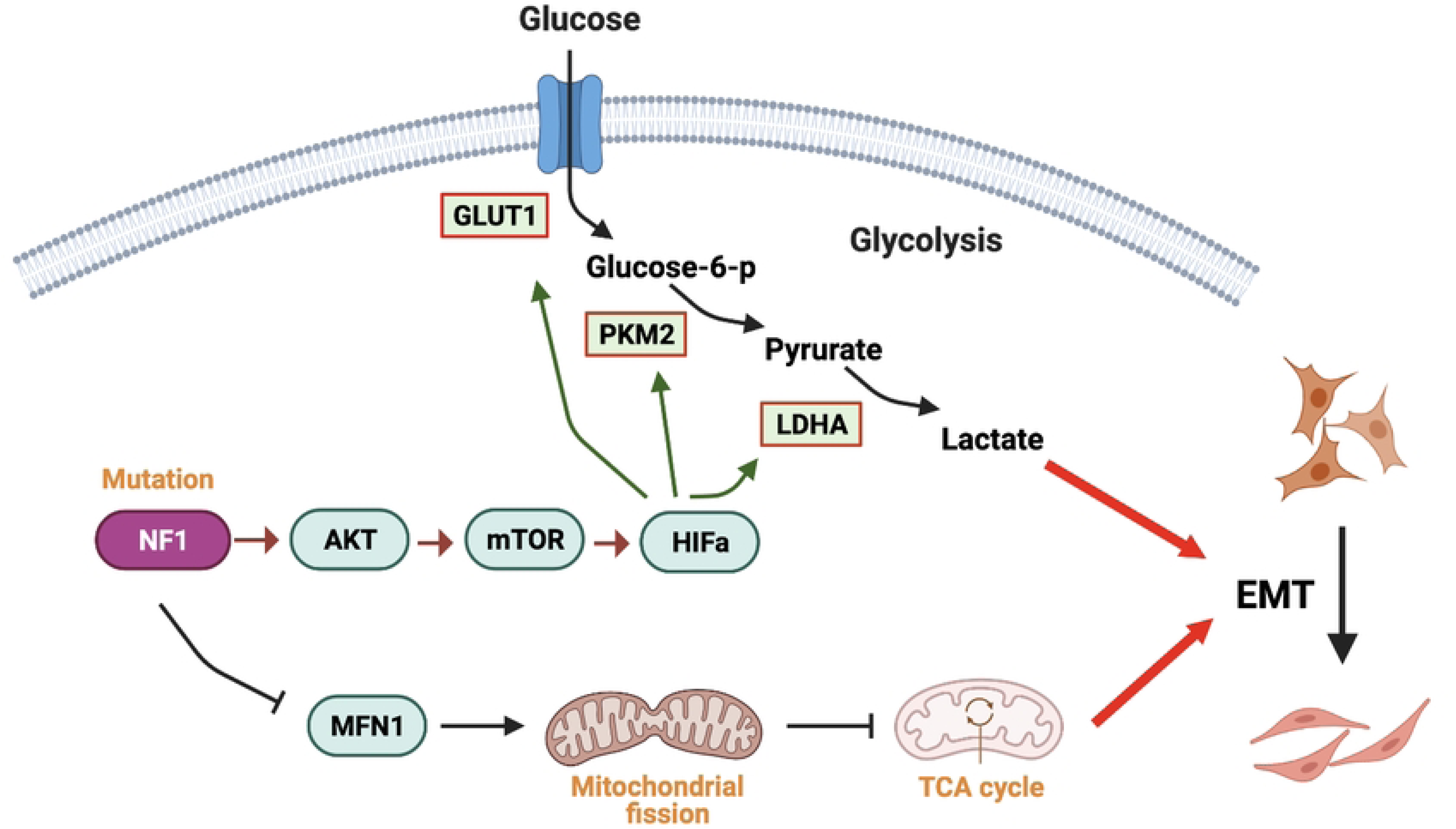
Molecular mechanism of NF1 mutation remodeling energy metabolism and promoting EMT of GBM cells.

## Acknowledgements

Not applicable.

## Author contributions

Y.W.P. and G.Q.Y. designed and supervised the experiments. Q.D., J.S., B.W. and L.N. performed the experiments and wrote the main manuscript text. H.Y., provided glioma tissues of glioma and brain trauma patients. X.Q.W., J.Q.D and Q.L. guided the operation of experiment and analyzed data. All authors critically revised and approved the final manuscript.

## Data availability

Not applicable.

## Declarations

Ethics approval and consent to participate

The present study was approved by The Medical Ethics Committee of The Second Hospital of Lanzhou University.

## Consent for publication

Not applicable.

